# FLOWERING LOCUS C integrates carbon and nitrogen signaling for the proper timing of flowering in Arabidopsis

**DOI:** 10.1101/2024.02.15.580522

**Authors:** Vladislav Gramma, Justyna Jadwiga Olas, Vasiliki Zacharaki, Jathish Ponnu, Urszula Marta Luzarowska, Magdalena Musialak-Lange, Vanessa Wahl

**Author notes:** Corresponding author/author responsible for distribution of materials. These three authors (in alphabetical order) contributed equally (Vladislav Gramma, Justyna Jadwiga Olas, Vasiliki Zacharaki). University of Applied Sciences Berlin (VG); Leibniz Institute of Vegetable and Ornamental Crop, Großbeeren, Germany (JJO); Metasysx GmbH, Am Mühlenberg 11, Potsdam, Germany (MML).

## Abstract

The timing of flowering in plants is modulated by both carbon (C) and nitrogen (N) signaling pathways. In a previous study, we established a pivotal role of the sucrose-signaling trehalose 6-phosphate pathway in regulating flowering under N-limited short-day conditions. In this work, we expand on our finding that wild-type plants grown under N-limited short days require an active trehalose 6-phosphate pathway to be able to flower. Both wild-type plants grown under N-limited conditions and knock-down plants of *TREHALOSE PHOSPHATE SYNTHASE1* induce *FLOWERING LOCUS C* expression, a well-known floral repressor associated with the vernalization response. When exposed to an extended period of cold, a mutant of *FLOWERING LOCUS C* fails to respond to N availability, and flowers at the same time under N-limited and full-nutrition conditions. Our data suggest that SUCROSE NON-FERMENTING 1 RELATED KINASE 1-dependent trehalose 6-phosphate-mediated C signaling and a novel mechanism downstream of N signaling likely involving NIN-LIKE PROTEIN 7 impact the expression of *FLOWERING LOCUS C.* Collectively, our data underscore the existence of a multi-factor regulatory system in which both C and N signaling pathways jointly govern the regulation of flowering in plants.

## Introduction

Owing to their sessile nature, plants adapt to environmental changes by modifying their development and growth. These processes require significant amounts of energy. Plants are in constant feedback with the environment and their nutrient status, especially carbon (C) and nitrogen (N), that serve as crucial bases for energy production and biomass generation. Low levels of C or N in the cells suppresses development and growth in plants and triggers the onset of senescence. To balance energy-intensive developmental processes with endogenous nutrient availability, plants have evolved intricate signaling networks (Fernie et al., 2020).

Flowering is an important developmental process in the life cycle of plants with correct timing being essential for reproductive success. It is regulated by a sophisticated genetic network that integrates various environmental and endogenous signals to regulate the expression of the floral integrator genes such as the florigen, *FLOWERING LOCUS T* (*FT*), and *SUPPRESSOR OF OVEREXPRESSION OF CONSTANS 1* (*SOC1*) (Srikanth and Schmid, 2011; Romera-Branchat et al., 2014; Song et al., 2015). FT integrates signals perceived in the leaves and conveys this information to the shoot apical meristem (SAM) to induce flowering (Corbesier et al., 2007; Jaeger and Wigge, 2007; Mathieu et al., 2007). At the SAM, FT interacts with the bZIP transcription factor FLOWERING LOCUS D (FD) to form a complex that directly activates *SOC1* along with floral meristem identity genes such as *APETALA1* (*AP1*) (Abe et al., 2005; Wigge et al., 2005).

In addition to other stimuli, temperature impacts greatly the time of flowering. Increased ambient temperature results in earlier flowering due to decreased SVP protein stability (Lee et al., 2013; Lee et al., 2014). SVP forms a temperature-dependent flowering repressor complex with partners such as FLOWERING LOCUS M (FLM)/MADS AFFECTING FLOWERING1 (MAF1), an orthologue of FLOWERING LOCUS C (FLC) (Pose et al., 2013; Sureshkumar et al., 2016), resulting in earlier flowering when plants are exposed to warmer conditions (Pose et al., 2013). SVP was also shown to interact with FLC in a flowering repressor complex (Fujiwara et al., 2008; Li et al., 2008). This delays floral transition by directly reducing the expression of *FT*, *FD,* and *SOC1* (Hepworth et al., 2002; Helliwell et al., 2006; Searle et al., 2006; Lee et al., 2007; Li et al., 2008). In winter-annual accessions of *Arabidopsis thaliana* (Arabidopsis) flowering is suppressed due to active *FRIGIDA* (*FRI*) resulting in promoted expression of *FLC*, unless the plants are exposed to a long period of cold (vernalization process) (Sheldon et al., 2000). This regulation involves a plethora of proteins and complexes acting in many layers of gene regulation, ranging from RNA structures, epigenetic modification to transcriptional and mRNA processing control (reviewed in Whittaker and Dean, 2017; and Sharma et al., 2020; Xu et al., 2021; Xu et al., 2021; Yang et al., 2022).

Organic C and N supply is essential in particular for vegetative growth and plant development (Sulpice et al., 2013). It is known that nutrients are essential for developmental transitions (Fernie et al., 2020), but the underlying mechanisms continue to be subject to active investigation. Interestingly, *FLC* expression was observed to increase significantly in NITRATE TRANSPORTER 1.1 (NRT1.1) and NRT1.13 defective mutant plants (Teng et al., 2019; Chen et al., 2021). While NRT1.13 is suggested to be a nitrate transporter, NRT1.1 is a key component of nitrate signaling functioning as both a transporter and a sensor in roots (Li et al., 2021). This suggests a nitrate signaling-dependent control of *FLC* as proposed by Kant and colleagues (Kant et al., 2011). This is supported by the introduction of an *flc-3* mutation into the late-flowering NRT1.1 deficient plant background which restored wild-type flowering (Teng et al., 2019).

Previous studies have identified multiple factors that influence N-regulated flowering, which often vary and depend on the cultivation systems used (Lin and Tsay, 2017). We are using a soil-based N-limited system developed by (Tschoep et al., 2009), which allows plant adaptation and the investigation of flowering time without stress-related symptoms (Olas et al., 2019; Olas et al., 2021). With this system we previously reported that nitrate-regulated flowering depends on SAM factors. Notably, in N-limiting conditions, nitrate-responsive gene expression is affected and nitrate assimilation is reduced in the SAM (Olas et al., 2019). The early nitrate response involves the NIN-LIKE PROTEIN (NLP) transcription factors NLP6 and NLP7. They accumulate in the nucleus in the presence of nitrate, regulating gene expression through nitrate responsive cis-elements (NRE) (Konishi and Yanagisawa, 2013; Marchive et al., 2013). Limited nitrate availability delays flowering due to decreased expression of *SOC1*, likely through NLP6/NLP7-regulated expression of the SQUAMOSA PROMOTER-BINDING PROTEIN-LIKE transcription factors encoding genes *SPL3* and *SPL5* (Olas et al., 2019).

The sucrose signal trehalose 6-phosphate (T6P) regulates a plethora of developmental and physiological responses (reviewed in Fichtner and Lunn, 2021). In Arabidopsis, T6P is synthesized by TREHALOSE PHOSPHATE SYNTHASE1 (TPS1) (Vandesteene et al., 2010; Yang et al., 2012) and it acts mainly by modulating the SUCROSE NON-FERMENTING 1 RELATED KINASE 1 (SnRK1) activity. Moreover, T6P was suggested to be able to bind directly to the SnRK1 upstream activating kinases and inhibit their activity (Zhai et al., 2018). SnRK1 is a key sensor of energy status and it is required for both normal growth and plant responses to stresses that impact plant fitness and survival (Polge and Thomas, 2007; Baena-Gonzalez and Sheen, 2008). Although single mutants of SnRK1 catalytic subunits resemble wild-type plants (Baena-Gonzalez et al., 2007; Jeong et al., 2015), *tps1* mutants (*tps1-2*) are embryo-lethal (Eastmond et al., 2002). This can be bypassed by ectopically expressing dexamethasone-inducible *TPS1* (*GVG::TPS1*) during seed set (van Dijken et al., 2004). However, plants grown from these seeds remain in the vegetative phase for a highly extended period or fail to flower entirely (van Dijken et al., 2004; Wahl et al., 2013). T6P signaling induces flowering in leaves via *FT* and also acts at the SAM through microRNA156 (miR156) and its target transcripts, *SPL3-5* (Wahl et al., 2013), at least partially via the modulation of the SUCROSE NON-FERMENTING 1 RELATED KINASE 1 (SnRK1) complex activity (Zacharaki et al., 2022). This was supported by the observation that loss of SnRK1 activity in the *tps1-2,GVG::TPS1* plants led to early induction of *FT* in the leaves, reduced *miR156* levels and strong induction of *SPL3* in the SAM during bolting (Zacharaki et al., 2022). Taken together these findings indicate that both C and N signaling can target the same components of the flowering network at the SAM (Wahl et al., 2013; Olas et al., 2019; Zacharaki et al., 2022), underscoring their joint importance for the proper timing of flowering.

Even though the current understanding implies a straightforward output downstream of nutrient signaling, our data now indicate a more complex relationship between nutrient signaling and developmental programs. Here, we demonstrate that the T6P pathway, which controls flowering under N limitation in short days (Olas et al., 2019), impacts on the expression of *FLC* in addition to *FLC* being differentially expressed upon exposure to contrasting N levels. Our findings suggest that both C- and N-dependent pathways regulate Arabidopsis flowering time by modulating *FLC* expression, implying a role in the composition and timing of the FLC-SVP repressor complex within a developmental context.

## Results

### Sucrose signaling represses *FLC*

We have previously reported that plants grown under N-limited conditions and short days (SD) accumulate both sucrose and T6P towards the end of the vegetative growth phase. Importantly, *TPS1* knock-down plants (*35S::amiRTPS1*) did not flower under these conditions (Olas et al., 2019). To understand this phenomenon, we analyzed a developmental series of rosette samples from both Col-0 and *35S::amiRTPS1* plants, focusing on candidate genes, which specifically change their expression before the floral transition. This analysis included multiple flowering time genes assessed by RT-qPCR. Notably, we found a strong up-regulation of *FLC* expression in 4- to 6-days-old and *MAF5* (*MADS AFFECTING FLOWERING 5*) expression in 12-days-old *35S::amiRTPS1* plants (Fig. 1A, Fig. S1), similar to Zeng et al. (2024). Considering that *FLC* expression has previously been suggested to be modulated in response to N availability (Kant et al., 2011), it is an interesting candidate for further investigation. We observed that *FLC* expression declines before the floral transition, which occurs at 10 days after germination (DAG) in Col-0 wild-type plants and 19 DAG in *35S::amiRTPS1* grown in full nutrition soil (Wahl et al., 2013). This suggests that the T6P pathway fundamentally contributes to the full repression of *FLC* in young seedlings. While we initially did not anticipate that the T6P pathway could affect *FLC* expression at later stages, we found that when the *flc-3* mutation is introduced into the *tps1-2 GVG::TPS1* background, it partially rescues the late flowering and the delayed vegetative phase transition observed in this *tps1-2 GVG::TPS1* (Fig. 1B,C; Fig. S2; Table S1,2). Our data, therefore, suggest that the T6P pathway is involved in *FLC* regulation to promote flowering and facilitate the vegetative phase change.

**Figure 1.**
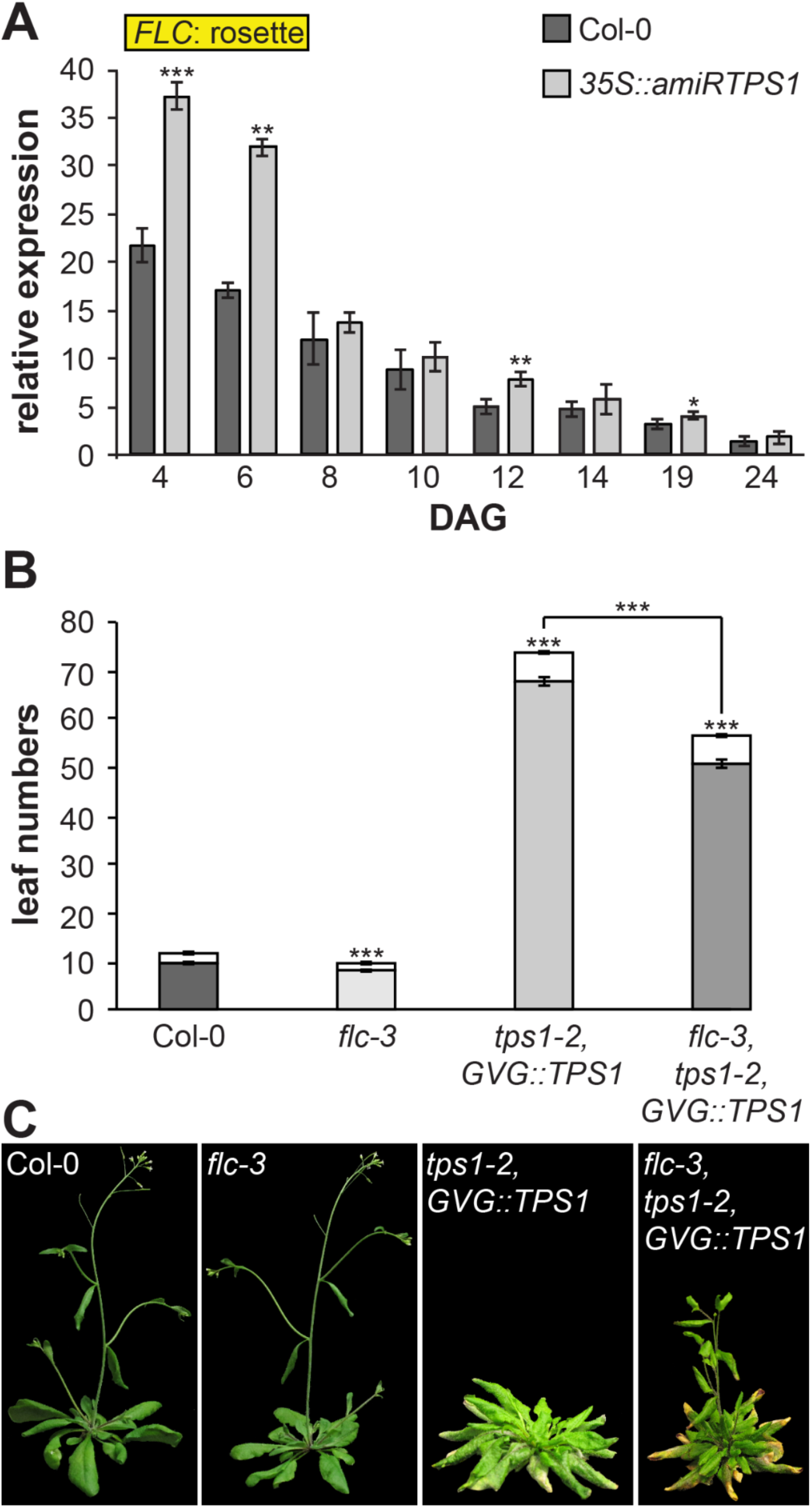
The Trehalose 6-phosphate pathway impacts on *FLOWERING LOCUS C*. (**A**) Expression of *FLOWERING LOCUS C* measured by RT-qPCR in rosettes of Col-0 and *35S::amiRTPS1* plants grown under long days (16h light/ 8h darkness). *n* = 4. (**B**) Flowering time measured as leaf numbers (rosette leaves in gray; cauline leaves in white). n ! 15 individual plants per genotype. (**C**) Representative photographs of the plants analyzed in (B). Abbreviations:days after germination (DAG). Data represents mean, error bars are standard deviations (s.d.), statistically significant difference compared to Col-0 wild-type (Student *t*-test, \**P*<0.05, \*\**P*<0.01, and \*\*\**P*<0.001).

### FLC integrates N-signaling into the flowering network

It has been previously shown that flowering is delayed in wild-type plants grown in the limited N (LN) soil (Olas et al., 2019). Furthermore, some data suggest that *FLC* expression may be influenced by N availability (Kant et al., 2011). Thus, we conducted experiments to investigate the potential regulation of *FLC* expression by N status. We grew wild-type Col-0 plants in a soil-based growth system (Tschoep et al., 2009), consisting of a soil with optimal N (ON) and one with LN source. We observed elevated *FLC* expression levels in LN in both rosettes and apices of Col-0 plants grown continuously in SD conditions (Fig. 2A, B), and in apices of plants that were initially grown in SD conditions and subsequently transferred to LD conditions (Fig. 2C). Similarly, we found upregulation of *MAF5* levels in rosettes of LN grown plants (Fig. S3). Considering that the *MAF5* was found to act downstream of the T6P pathway, this finding suggests a link between N and T6P signaling pathways. (Fig. S1), *MAF5* was also found to be upregulated in response to N limitation (Fig. S3). Next, to obtain information on the expression pattern at higher spatial resolution, we used *FLC* as a probe and performed RNA *in situ* hybridization (Fig. 2D). *FLC* transcript was detectable at the SAM and in young leaves of LN grown plants, confirming our previous observations that limited N availability enhances *FLC* expression in plants. This finding suggests that FLC plays a role in the regulation of flowering time in response to N availability.

**Figure 2.**
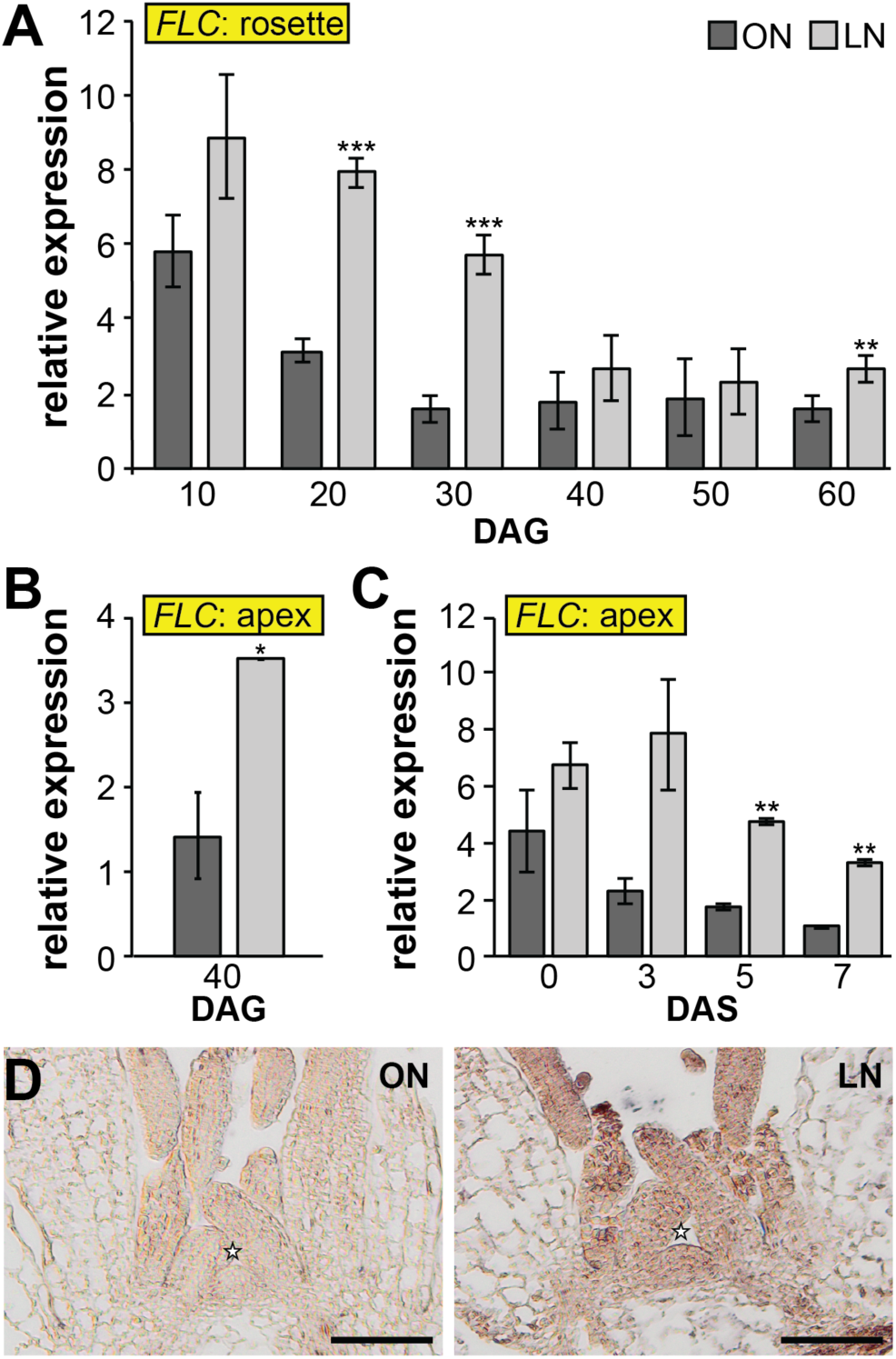
*FLOWERING LOCUS C* in response to nitrogen limitation. (**A, B**) Expression of *FLOWERING LOCUS C* measured by RT-qPCR in rosettes (A) and apices (B) of Col-0 plants grown in optimal nitrogen (ON) and limited-nitrogen (LN) conditions under short days (16h light/ 8h dark). (**C**) *FLC* expression measured by RT-qPCR in apices of plants initially grown under short days (30 days) and then transferred to long days to initiate the floral transition for 3, 5, and 7 days. (**D**) RNA *in situ* hybridization using *FLOWERING LOCUS C* specific probe on longitudinal sections through vegetative apices of Col-0 plants grown in ON and LN soils. Abbreviations: days after germination (DAG); days after shift (DAS). Data represents mean, error bars are standard deviations (s.d.), n=3, statistically significant difference between ON and LN (Student’s *t*-test, \**P*<0.05, \*\**P*<0.01, \*\*\**P*<0.001). Star indicates apex summit.

It is well established that exposure to low temperatures decreases *FLC* expression in plants (Searle et al., 2006). For this reason, we grew wild-type plants at 4°C in SD for 8 weeks, followed by a transfer to 22°C until flowering. This treatment resulted in wild-type plants flowering at the same time in both N regimes, suggesting that FLC contributes to the delayed flowering time observed in plants grown in the LN soil (Fig. 3A; Table S1).

**Figure 3.**
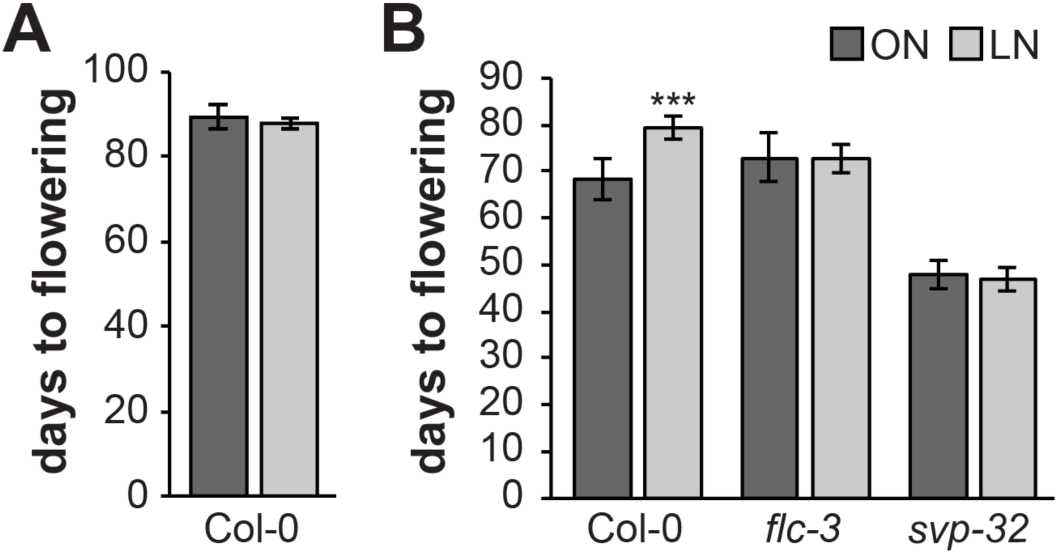
FLOWERING LOCUS C and SHORT VEGETATIVE PHASE are required for the limited nitrogen-dependent flowering response. (**A**) Flowering time of Col-0 wild-type plants treated with an 8-week period of cold. Note that afterwards plants were transferred to 22°C until flowering. (**B**) Flowering time of Col-0, *flc-3,* and *svp-32* mutant plants grown under short-day (8h light/16h darkness) conditions. Data represents mean, error bars are standard deviations (s.d.), n ! 15 individual plants per genotype, statistically significant difference between ON and LN (Student’s *t*-test, \*\*\**P*<0.001).

FLC is known to form a flowering repressor complex with SVP to suppress *SOC1* at the SAM (Li et al., 2008). Unlike *FLC*, *SVP* was not differentially expressed in either LN-grown plants or *TPS1* knock-down plants (Fig. S4; Fig. S5). Importantly, neither *flc-3* nor *svp-32* mutant plants responded to the reduced N content in the LN soil (Fig. 3B; Table S1), flowering at the same time in ON and LN conditions. This indicates that both FLC and SVP play a role in the N-dependent regulation of flowering time.

### N-signaling affects *FLC* via NLP7

NLPs are key regulators of nitrate sensing and signaling, with NLP6 and NLP7 being two of the most well-characterized members of this family in *Arabidopsis* (Fredes et al., 2019). In the presence of nitrate, NLP7 is retained in the nucleus through phosphorylation, where it binds to NREs present in N-responsive genes to promote their expression (Konishi and Yanagisawa, 2013). Interestingly, we observed a significant reduction of *FLC* expression in the *nlp7-1* mutant, indicating that an active NLP7 modulates *FLC* expression when N is not limited (Fig. 4).

**Figure 4.**
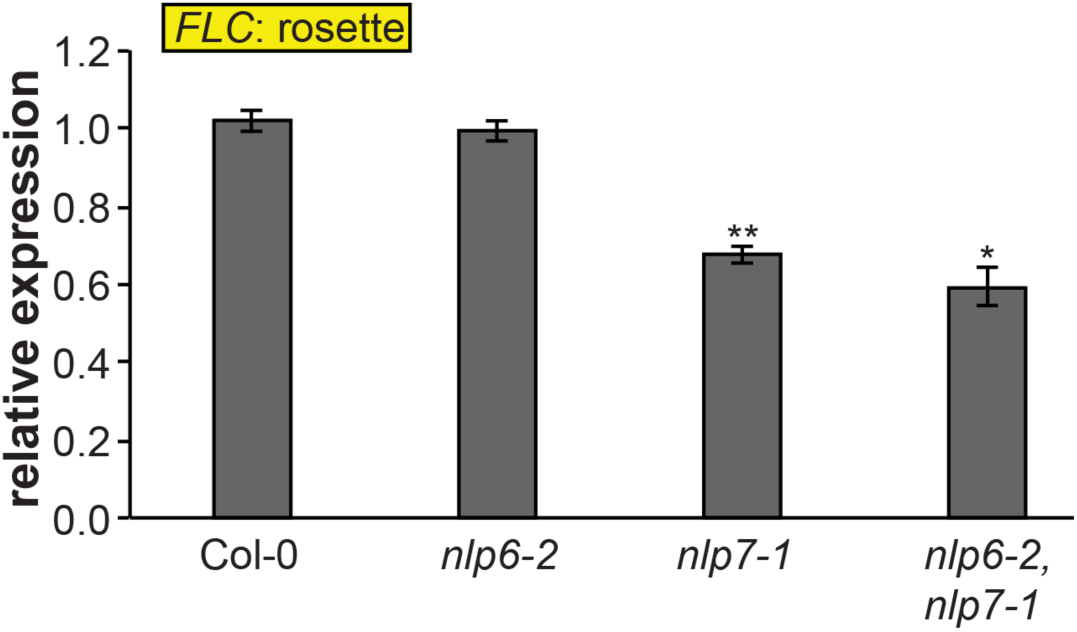
*FLOWERING LOCUS C* expression downstream of NIN-LIKE PROTEIN 6 (NLP6) and NIN-LIKE PROTEIN 7 (NLP7). Expression of *FLOWERING LOCUS C* measured by RT-qPCR at 10 days after germination (DAG) in rosettes of Col-0, *nlp6-2*, *nlp7-1*, and *nlp6-2,nlp7-1* plants. Data represents mean, error bars are standard deviations (s.d.), n=3, statistically significant difference compared to Col-0 wild-type (Student’s *t*-test, \**P*<0.05, \*\**P*<0.01).

Since the *FLC* gene does not carry an NRE in its promoter, genomic or downstream sequences, we expanded our analysis to include other flowering time genes that regulate *FLC* (Table S3). Notably, *FRI*, a key regulator upstream of *FLC,* has four putative NREs (Table S3). However, in the Col-0 background, the *FRI* locus encodes an inactive protein and therefore does not influence *FLC*. In addition to *FRI*, other genomic loci encoding *FLC* regulators were also found with putative NREs (Table S3), but their expression was unaffected under N limited conditions (Fig. S6). This was also the case in *35S::amiRTPS1* plants (Fig. S7). Taken together, this suggests that *FLC* suppression involves an as yet unknown transcription factor(s), whose activity is regulated by NLP7.

### Sucrose and N-signals interconnect at the level of *FLC* for coordinated flowering time regulation

We have previously demonstrated that the T6P pathway and sufficient nitrate levels are necessary for floral induction in SD (Olas et al., 2019). The fact that *35S::amiRTPS1* plants fail to flower when N is limited and that *FLC* expression is modulated by N availability, prompted us to test whether *FLC* is a target of both N signaling and the T6P pathway.

We observed that *FLC* transcription was elevated in rosettes of wild-type plants grown under SD with limited N which was even more pronounced in *35S::amiRTPS1* plants (Fig. 5A). This suggests an additive effect between N signaling and the T6P pathway, both converging on the *SPL3-5* node at the SAM (Wahl et al., 2013; Olas et al., 2019). To test whether *FLC* could be regulated through SPL3-5, we measured its expression in *spl345* mutants (Xie et al., 2020). However, *FLC* expression in rosette leaves of *spl345* mutants was comparable to that of wild-type plants (Fig. S8A), indicating that both pathways regulate *FLC* expression via another mechanism. Similarly to *FLC*, we did not observe any difference in *SVP* expression in *spl345* compared to Col-0 plants (Fig. S8B).

**Figure 5.**
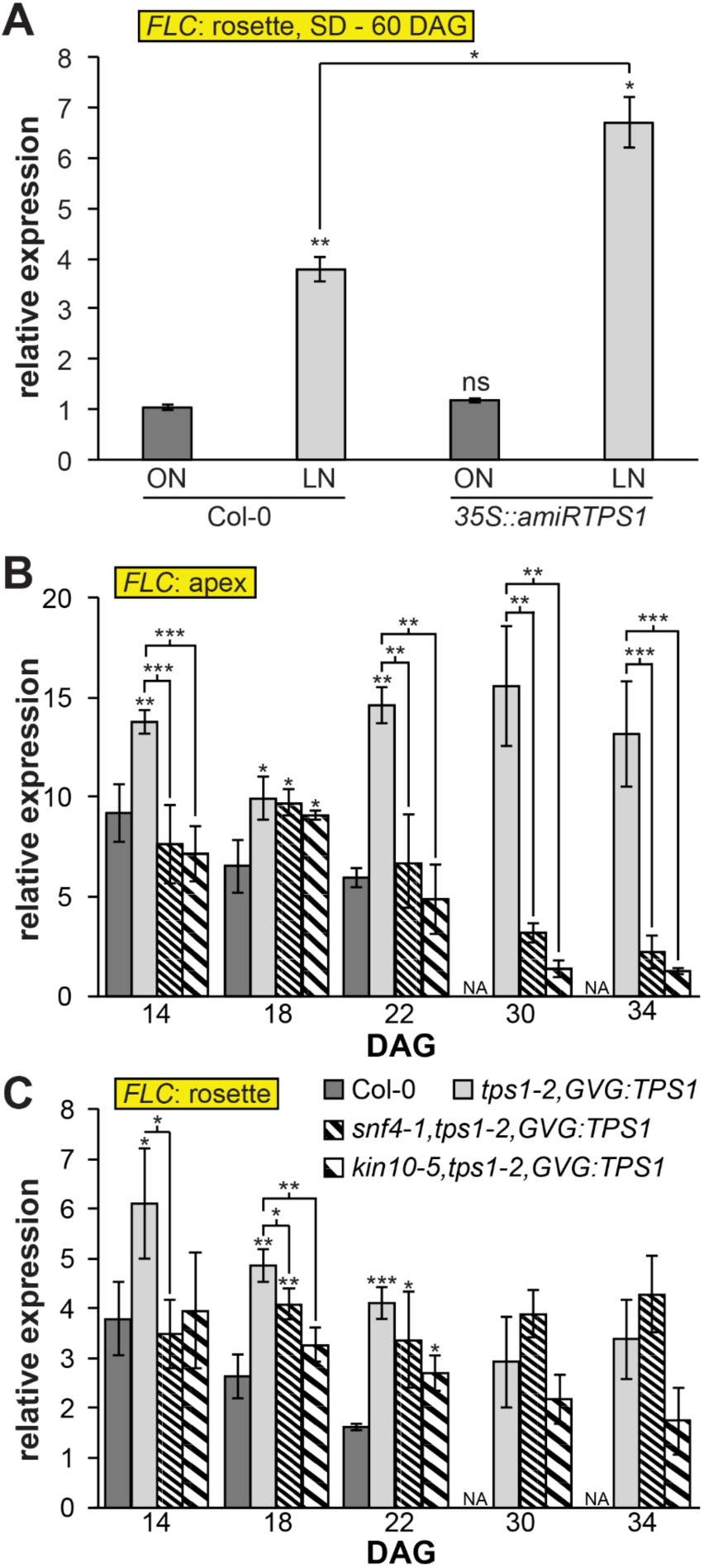
Trehalose 6-phosphate pathway and nitrogen-signaling converge at *FLOWERING LOCUS C*. (**A**, **B**, **C**) Expression of *FLOWERING LOCUS C* measured by RT-qPCR in (A) rosettes of wild-type Col-0 and *35S::amiRTPS1* plants grown in optimal nitrogen (ON) and limited-nitrogen (LN) conditions under short days (8h light/ 16h dark) at 60 days after germination (DAG), in (B) apices and (C) rosettes of wild-type Col-0, *tps1-2,GVG:TPS1*, *snf4,tps1-2,GVG:TPS1* and *kin10,tps1-2,GVG:TPS1* plants grown in standard soil under long days (16h light/ 8h dark). Data represents mean, error bars are standard deviations (s.d.), significant difference compared to Col-0 wild-type (Student *t*-test, \**P*<0.05, \*\**P*<0.01, and \*\*\**P*<0.001).

The T6P pathway is known to function by directly modulating SnRK1 activity (Zhang et al., 2009). Loss of SnRK1 activity restores flowering of *tps1* mutants in LD by initial induction of *FT* in the leaves and subsequent suppression of miR156 followed by SPLs induction in the SAM (Zacharaki et al., 2022). Thus, we tested whether *FLC* regulation in *tps1* mutants is also mediated by SnRK1. We found that indeed *FLC* expression was increased in the *tps1 GVG::TPS1* mutant where SnRK1 is fully active. Interestingly, introducing non-catalytically active mutations in SnRK1 within the *tps1 GVG::TPS1* background restores *FLC* expression to wild-type levels in both rosette leaves and apex tissue (Fig. 5B,C). The suppression of *FLC* in the double mutant is more pronounced in the apex than the rosette leaves, underscoring the critical role of the T6P pathway in controlling developmental transitions. Our data suggest that *FLC* expression is regulated by both nitrate and sugar availability via NPLs and the T6P pathway through the SnRK1 complex, respectively (Fig. 6).

**Figure 6.**
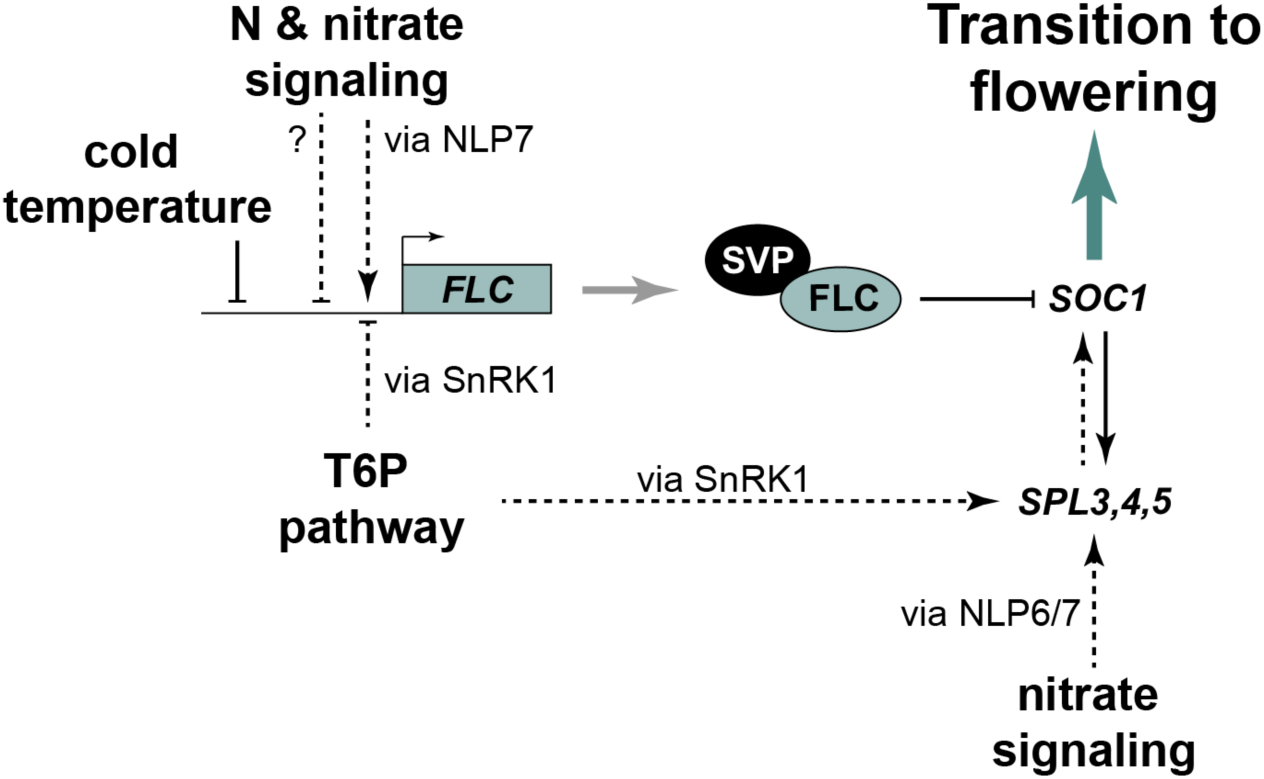
Carbon and nitrogen signaling target similar components of the flowering network in the shoot apical meristem for the proper timing of flowering. *FLOWERING LOCUS C,* a key repressor of flowering, is not only regulated by cold temperature as part of the vernalization process, but is also affected by nutrient availability. The Trehalose 6-phosphate pathway negatively impacts *FLOWERING LOCUS C* via SUCROSE NON-FERMENTING 1 RELATED KINASE 1. Nitrogen signaling controls *FLOWERING LOCUS C* via a yet to identify mechanism involving NIN-LIKE PROTEIN 7. The repressor complex composed of FLOWERING LOCUS C and SVP is eventually tuned by the adjustment of *FLOWERING LOCUS C* expression downstream of both carbon and nitrogen signaling to control *SUPPRESSOR OF OVEREXPRESSION OF CONSTANS 1* in the shoot apical meristem. Independently, both C and N pathways work via the age pathway (SQUAMOSA PROMOTER-BINDING PROTEIN-LIKE 3-5) to induce flowering.

## Discussion

C and N are essential for plant growth and development and the ability of plants to properly sense their availability is crucial due to their sessile nature. C in the form of sucrose is produced via photosynthesis in the leaves while N can be taken up in both inorganic forms, as nitrate and ammonia, or organic forms, as amino acids.

In *Arabidopsis*, a key sugar sensor is the T6P pathway which functions via SnRK1 activity. The T6P pathway has a key role in plants’ developmental transitions, such as flowering. So far, it has been shown that both *miR156* and *FT* regulation in the SAM and leaves, respectively, are required for *tps1* plants to complete their transition to flowering (Wahl et al., 2013; Ponnu et al., 2020). Here we found that *FLC,* a repressor of flowering, is also regulated by the T6P pathway (Fig. 1A) and that loss of functional *FLC* partially restores flowering in *tps1* (Fig. 1B,C). Although, we do not expect that *FLC* regulation is the prime target of the T6P pathway under normal growth conditions, it could represent an additional mechanism to prevent flowering under non-optimal growth conditions.

Plants experiencing a sudden shift to colder temperatures have increased amounts sucrose previously proposed to serve as a freezing protectant and concomitant rising T6P levels (reviewed in Stitt and Hurry, 2002; Carillo et al., 2013). During long cold exposure, *FLC* is suppressed and this regulation involves several mechanisms, ranging from RNA structures to epigenetic control. *FLC* suppression allows induction of *FT* and *SOC1* and flowering to commence (Whittaker and Dean, 2017). In this scenario, when plants experience cooler temperatures, nutrients that provide plants with C and N, are transported and stored to serve as a basis for rapid growth for when conditions are optimal again or used and metabolically transformed into cryoprotectants to protect the cells from freezing damage (Kaplan et al., 2007). Thus, in sub-optimal growth conditions, the T6P pathway might contribute to the suppression of *FLC* in response to the C status.

N availability is a key factor in the regulation of plants’ developmental processes and phase transitions including the timing of flowering (Klebs, 1913; Dickens and Van Staden, 1988; Bernier et al., 1993; Olas et al., 2019). Arabidopsis cultivated on synthetic substrates exhibit early flowering in response to reduced N levels (Castro Marin et al., 2011; Kant et al., 2011; Liu et al., 2013). Conversely, soil-grown plants subjected to N limitation flower later than those cultivated in soil without N limitation, which we previously linked to the induction of *SPL3* and *SPL5* by NLP6 and NLP7 (Olas et al., 2019). In this study we discovered that this phenotype can additionally be explained by significantly elevated levels of *FLC* under N limitation (Fig. 2A-D). Furthermore, we found that flowering time in plants with suppressed *FLC* due to the vernalization response or with a non-functional *flc-3* allele is independent of N availability (Fig. 3A,B). These results demonstrate that despite the general belief that *FLC* does not play a major role in the regulation of flowering time in rapid-cycling accessions, such as Col-0, *FLC* is required for fine-tuning the timing of floral transition downstream of N signaling. Similar to *flc-3*, *svp-32* mutants flower at the same time in the ON and LN soils (Fig. 3B), suggesting a role of *SVP* in N-dependent flowering time regulation. However, in contrast to *FLC*, *SVP* is not differentially expressed in plants grown in ON and LN soil (Fig. S5). FLC and SVP proteins form a flowering repressor complex that delays floral transition by directly reducing the expression of *FT* and *SOC1* (Hepworth et al., 2002; Helliwell et al., 2006; Lee et al., 2007; Li et al., 2008). Given that both functional *FLC* and *SVP* loci are required for the adjustment of flowering time in response to N availability, it is likely that the N signal is integrated at the level of the FLC-SVP complex. In this scenario, the formation of the repressor complex would be tuned by the adjustment of *FLC* expression downstream of N-signaling. Several transcription factors that are transcriptionally responsive to the N status have been identified as prime responsive genes to N availability (Vidal et al., 2015). NLPs are transcription factors facilitating nitrate signaling in plants, with NLP6 and NLP7 representing the master regulators and the two most studied (Fredes et al., 2019). In the absence of nitrate, NLP7 localizes strictly to the cytosol, while exposure to nitrate triggers its localization into the nucleus where it binds directly to NREs of nitrate-regulated genes (Konishi and Yanagisawa, 2013; Marchive et al., 2013). Since NREs are not present in the *FLC* locus (Table S3), it is unlikely to be directly controlled by NLPs. Other examples of *FLC* regulation related to N availability, are the *nrt1.1* and *nrt1.13*, mutants of the nitrate sensor and transporter NRT1.1 and transporter NRT1.13 (Teng et al., 2019; Chen et al., 2021). Similar to our findings (Fig. S7), expression of known upstream regulators of *FLC* was not changed in *nrt1.13*, suggesting that NRT1.13 regulates *FLC* expression and flowering time independently of these known pathways (Chen et al., 2021).

Interestingly, we found that *FLC* was significantly downregulated in the late flowering *nlp7-1* and *nlp6-2 nlp7-1* mutants grown on standard soil (Fig. 4A, Fig. S9) indicating that NLP7 plays a role in the modulation of *FLC* expression. Given the fact that NLP7 was found to control most of the nitrate-related gene response (Marchive et al., 2013; Alvarez et al., 2020), the *nlp7-1* mutant is thought to mimic a low nitrate state. Hence, this result appears to contradict our observation of *FLC* accumulation in LN-grown plants (Fig. 2A,B). This could be explained by the presence of an unknown NLP-independent mechanism responsible for *FLC* upregulation in LN conditions. However, it should be noted that in contrast to the mutant background, functional NLP7 is still present in wild-type plants exposed to limited N. Thus, *nlp7-1* might not entirely mimic the low-nitrate state after all and the absence of a functional NLP7 likely leads to compensation by other NLPs. Furthermore, NLP proteins contain a PB1 domain, which mediates protein-protein interactions influencing NLP activity (Konishi and Yanagisawa, 2019). Given this, NLP7 might form a complex with an unknown *FLC* repressor, thereby preventing its nuclear localization under low-nitrate conditions. In the absence of NLP7 or when plants are grown under optimal N conditions, this potential repressor would localize into the nucleus, leading to a repression of *FLC* expression. It will be interesting to further dissect the mechanisms of *FLC* regulation downstream of N-signaling in the future.

Our data demonstrate that both the T6P and N-signaling pathways possibly affect *FLC* expression via different mechanisms. Previous studies have demonstrated that both pathways act via the miR156/SPLs node (Wahl et al., 2013; Olas et al., 2019; Ponnu et al., 2020; Zacharaki et al., 2022). In particular, the expression of *SPL3* and *SPL5* is reduced in plants grown in N-limited environment (Bi et al., 2007; Pant et al., 2009; Krapp et al., 2011; Liang et al., 2012; Fischer et al., 2013), suggesting a role for the miR156/SPL3/5 module in the regulation of flowering time when N is limited. Similarly, the T6P pathway acts via miR156 downregulation and SPL3-5 upregulation to induce flowering and the vegetative phase change (Wahl et al., 2013; Ponnu et al., 2020; Zacharaki et al., 2022). Although both pathways converge on the miR156/SPLs module, *FLC* regulation seems to be independent (Fig. S8).

T6P has a key role in promoting growth and development by suppressing SnRK1 complex activity, via direct binding to the SnRK1 upstream kinases (Zhai et al., 2018). In a previous study, it was shown that *FT* was induced in the double *tps1-2 GVG::TPS1 kin10-5* and *tps1-2 GVG::TPS1 snf4* mutant as early as in wild-type plants (Zacharaki et al., 2022). Although this early *FT* induction promoted the floral transition in wild-type plants within a few days, this was not the case in both double mutants. The elevated expression of *FLC* in rosette leaves of these mutants (Fig. 5C) could thus at least partially explain this phenomenon. *FLC* downregulation is directly correlated with early *FT* upregulation previously observed in the double mutants (Zacharaki et al., 2022). Interestingly, we observed that *FLC* was also downregulated in the double mutants in the apex (Fig. 5B) with more striking differences later on, coinciding with the timing of floral transition (Zacharaki et al., 2022). In addition, ectopic *FLC* expression in the SAM has been associated with delayed flowering and reduced *SOC1* and *FD* expression (Sheldon et al., 2002; Noh and Amasino, 2003; Searle et al., 2006). This is also the case in *tps1-2 GVG::TPS1*, while gene expression is restored in the double mutants (Fig. S10) (Zacharaki et al., 2022). Our data combined with the findings of Zeng et al. (2024) suggest that the regulation of SnRK1 activity is essential for T6P-dependent floral induction, which has several modes of action throughout the floral network to ensure that sufficient energy is available for this demanding developmental transition. Finally, our findings shed further light on the multifactorial aspects of C- and N-dependent regulation of flowering time.

## Material and Methods

### Plant material and growth conditions

*Arabidopsis thaliana* plants used for this study are of the Columbia (Col-0) ecotype. Mutant and transgenic lines such as *flc-3, svp-32*, *35S::amiRTPS1*, *tps1-2,GVG::TPS1*, *tps1-2,GVG::TPS1,kin10-5*, *tps1-2,GVG::TPS1,snf4-1*, *nlp6-2*, *nlp7-1*, *nlp6-2,nlp7-1* and *spl345* were previously described (Michaels and Amasino, 1999; Lee et al., 2007; Wahl et al., 2013; Olas et al., 2019; Xie et al., 2020; Zacharaki et al., 2022). The *flc-3,tps1-2*,*GVG*::*TPS1* double mutant lines were generated by crossing. Genotypes were confirmed by a genotyping PCR using the oligonucleotides listed in Table S4.

Arabidopsis plants were grown in controlled growth chambers (Model E-36L, Percival Scientific Inc., Perry, IA, USA) at 22°C in long-day (LD, 16h light/8h dark) or short-day (SD, 8h dark/16h light) conditions. Light intensity was approximately 160 μmol/m²s. Controlled induction of flowering was performed by transferring the plants from non-inductive (SD) to inductive conditions (LD) as described (Schmid et al., 2003).

A previously established, almost natural, soil-based N-limited growth system consisting of ON and LN soil was used to grow plants (Tschoep et al., 2009). Briefly, the growth system consists of two types of peat-based soil mixtures with either an optimal level of N (ON, ∼850 mg (N)/kg) or a limited level of N (LN, ∼40 mg (N)/kg). Soil mixtures were prepared as described (Olas et al., 2019).

### Phenotypic analyses

Flowering time was defined as days to flowering (DTF), which describes the days after germination to the day of bolting (inflorescence length 0.5cm), and by the total number of leaves (TLN). At least 16 plants were used to determine flowering time of each genotype. For vegetative phase change, juvenile leaf numbers were recorded and the leaf shape was digitally documented as described (Ponnu et al., 2020). A student’s *t*-test was used to test the significance of the phenotypic differences.

### Reverse transcription quantitative PCR (RT-qPCR)

Sampling, RNA extraction and RT-qPCR analysis of *FLC* in the *tps1-2,GVG::TPS1*, *tps1-2,GVG::TPS1,kin10-5* and *tps1-2,GVG::TPS1,snf4-1* were performed as described (Zacharaki et al., 2022). RNA extraction and RT-qPCR analyses of all the other genes were performed according to Wahl et al. (2013). Relative expression values were calculated with the 2^DDCt method using Ct values of a housekeeping gene index of *TUB2* (At5g62690), *SAND* (At2g28390), *UBQ10* (At4g05320), and *PDF2* (At1g13320). RT-qPCR analyses were performed in three or four biological replicates (n=3 or 4). A Student’s *t*-test was used to test for statistical significance.

### RNA *in situ* hybridization

Wax embedding, sectioning, RNA *in situ* hybridization, and imaging were performed as described (Wahl et al., 2013; Gramma and Wahl, 2023). Probes were synthesized using the DIG RNA Labeling Kit (Roche, Mannheim, Germany) for CDS of the *FLC* gene cloned into the pGEM®-T Easy vector (Promega, Madison, Wisconsin, US). Oligonucleotides and construct IDs are listed in Table S4.

## Accession numbers

TPS1 (At1g78580), FLC (At5g10140), SVP (At2g22540), MAF5 (At5g65080), FCA (At2g19520), EMF1 (At5g11530), PIE1/SNF2 (At3g12810), NLP6 (At1g64530), NLP7 (At4g24020), FRI (At4g00650), SUF4 (At1g30970), ELF7 (At1g79730), SEF (At5g37055), VRN1 (At3g18990), VRN2 (At4g16845), EMF2/CYR1 (At5g51230), TFL2 (At5g17690), FVE (At2g19520), HUA2 (At2g19520), SNF4 (At1g09020), KIN10 (At3g01090), SPL3 (At2g33810), SPL4 (At1g53160), SPL5 (At3g15270).

## Data availability

The data supporting the findings of this study are included in this manuscript or the supplemental information and material can be obtained from the corresponding author upon reasonable request.

## Acknowledgments and Funding

We thank Rob Hancock, Markus Schmid and the Wahl group for discussions and comments on the manuscript, Armin Schlereth, Christin Abel, Kerstin Zander, and Sebastian Kindermann for technical support. Work in the Wahl group was supported by the BMBF (031B0191), the DFG (SPP1530: WA3639/1-2, 2-1), the Max Planck Society and the James Hutton Institute.

## Author’s contribution

VW conceived and designed the experiments and prepared the figures. All authors performed essential experiments and analyzed data: JJO and VW performed the RNA *in situ* hybridizations; VG, JJO, VZ, and MML the RT-qPCRs; JJO, UML and JP performed phenotypic analyses. VG, VZ and VW wrote the manuscript with contributions from the other authors. All authors have read and commented on the text and figures within this manuscript.

## Supplemental Material

**Supplemental Figure S1.** MADS AFFECTING FLOWERING 5 in *35S::amiRTPS1* plants.

**Supplemental Figure S2.** *flc-3* partially suppresses the delayed vegetative phase change phenotype of *tps1-2*,*GVG::TPS1* plants.

**Supplemental Figure S3.** *MADS AFFECTING FLOWERING 5* in response to nitrogen limitation.

**Supplemental Figure S4.** SHORT VEGETATIVE PHASE in *35S::amiRTPS1* plants.

**Supplemental Figure S5.** *SHORT VEGETATIVE PHASE* in response to nitrogen limitation.

**Supplemental Figure S6.** Regulators upstream *FLOWERING LOCUS C* in response to N limitation.

**Supplemental Figure S7.** Regulators upstream *FLOWERING LOCUS C* in *35S::amiRTPS1* plants.

**Supplemental Figure S8.** FLOWERING LOCUS C and *SHORT VEGETATIVE PHASE* in *spl345* mutant plants.

**Supplemental Figure S9.** Plant phenotype of *nlp6* and *nlp7* mutant plants.

**Supplemental Figure S10.** FLOWERING LOCUS D in *snrk1,tps1-2,GVG::TPS1* mutants.

**Supplemental Table S1.** Flowering time data of experiments described in this study (Figure S2).

**Supplemental Table S2.** Vegetative phase change data of experiment described in this study

**Supplemental Table S3.** Putative nitrate responsive *cis*-elements (NREs) in regulators upstream *FLOWERING LOCUS C*.

**Supplemental Table S4.** Oligonucleotides used in this study.

