## Supplemental Data for "FLOWERING LOCUS C integrates carbon and nitrogen signaling for the proper timing of flowering in Arabidopsis"

**Author emails and ORCID**:

**Short title:** C and N signaling regulate FLC.

### Supplementary Figures

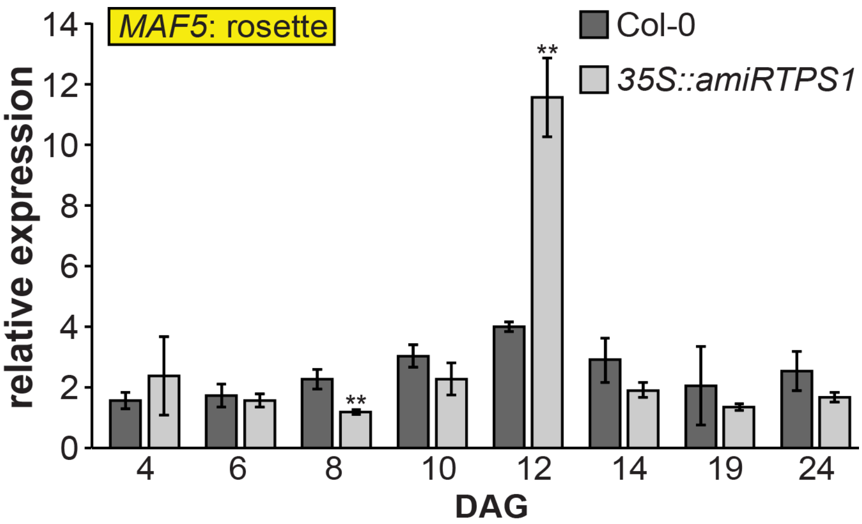


**Figure S1**: ***MADS AFFECTING FLOWERING 5* in *35S::amiRTPS1* plants.** Expression measured by RT-qPCR in rosettes of Col-0 and *35S::amiRTPS1* plants grown under long days (16h light/ 8h darkness). $n$ = 4. Data represents mean, error bars are standard deviations (s.d.), significant difference compared to Col-0 wild-type (Student $t$-test, $^{**}P<0.01$). Abbreviations: days after germination (DAG).

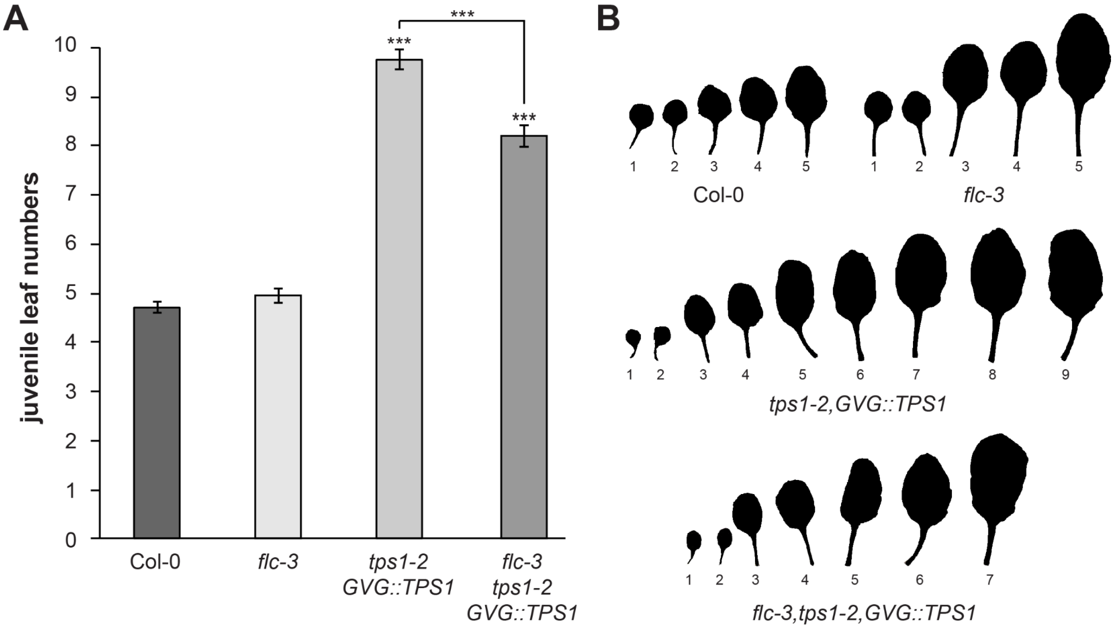


**Figure S2**: ***flc-3* partially suppresses the delayed vegetative phase change phenotype of *tps1-2,GVG::TPS1* plants.** (**A**) Number of juvenile leaves recorded from wild-type Col-0 *flc-3*, *tps1-2,GVG::TPS1*, and *flc-3,tps1-2,GVG::TPS1* plants grown under long days (16h light/ 8h darkness). $n$ = 20. (**B**) Leaf imprints of representative plants analyzed in (A). Data represents mean, error bars are standard deviations (s.d.), statistically significant difference compared to Col-0 wild-type (Student $t$-test, ***$P$<0.001).

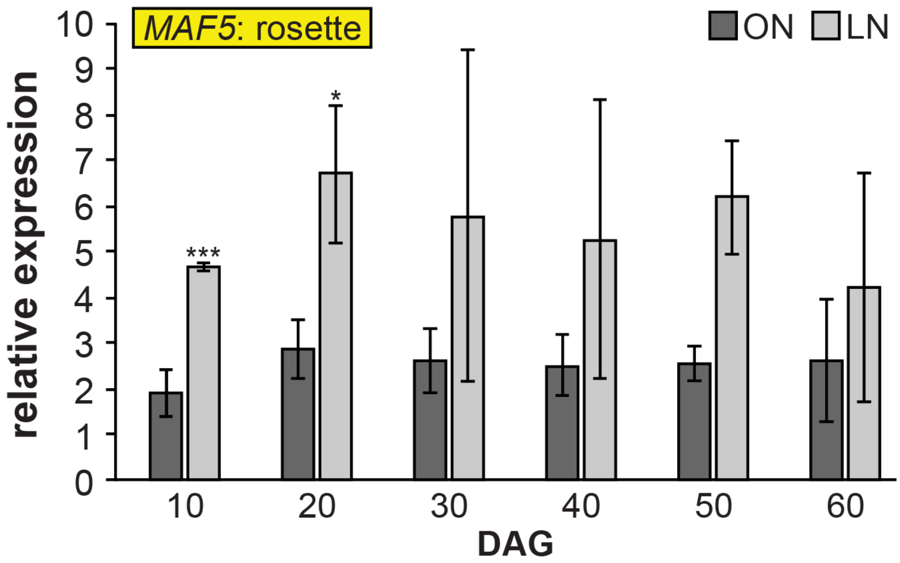


**Figure S3**: ***MADS AFFECTING FLOWERING 5* in response to nitrogen limitation.** Expression measured by RT-qPCR in rosettes of Col-0 plants grown in optimal nitrogen (ON) and limited-nitrogen (LN) conditions under short days (16h light/ 8h dark). $n = 3$. Data represents mean, error bars are standard deviations (s.d.), statistically significant difference between ON and LN (Student's *t*-test, *$P$<0.05, ***$P$<0.001). Abbreviations: days after germination (DAG).

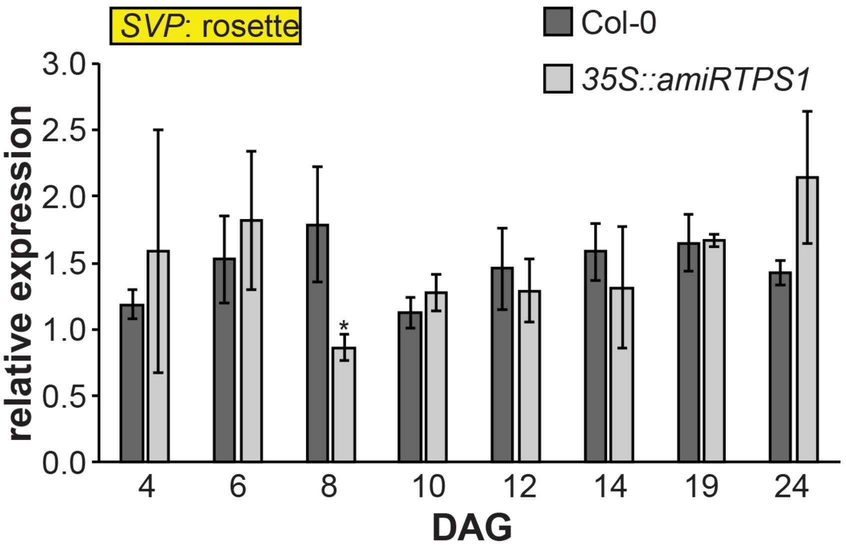


**Figure S4**: ***SHORT VEGETATIVE PHASE* in *35S::amiRTPS1* plants.** Expression measured by RT-qPCR in rosettes of Col-0 and *35S::amiRTPS1* plants grown under long days (16h light/ 8h darkness). *n* = 4. Data represents mean, error bars are standard deviations (s.d.), significant difference compared to Col-0 wild-type (Student *t*-test, **P*<0.05). Abbreviations: days after germination (DAG).

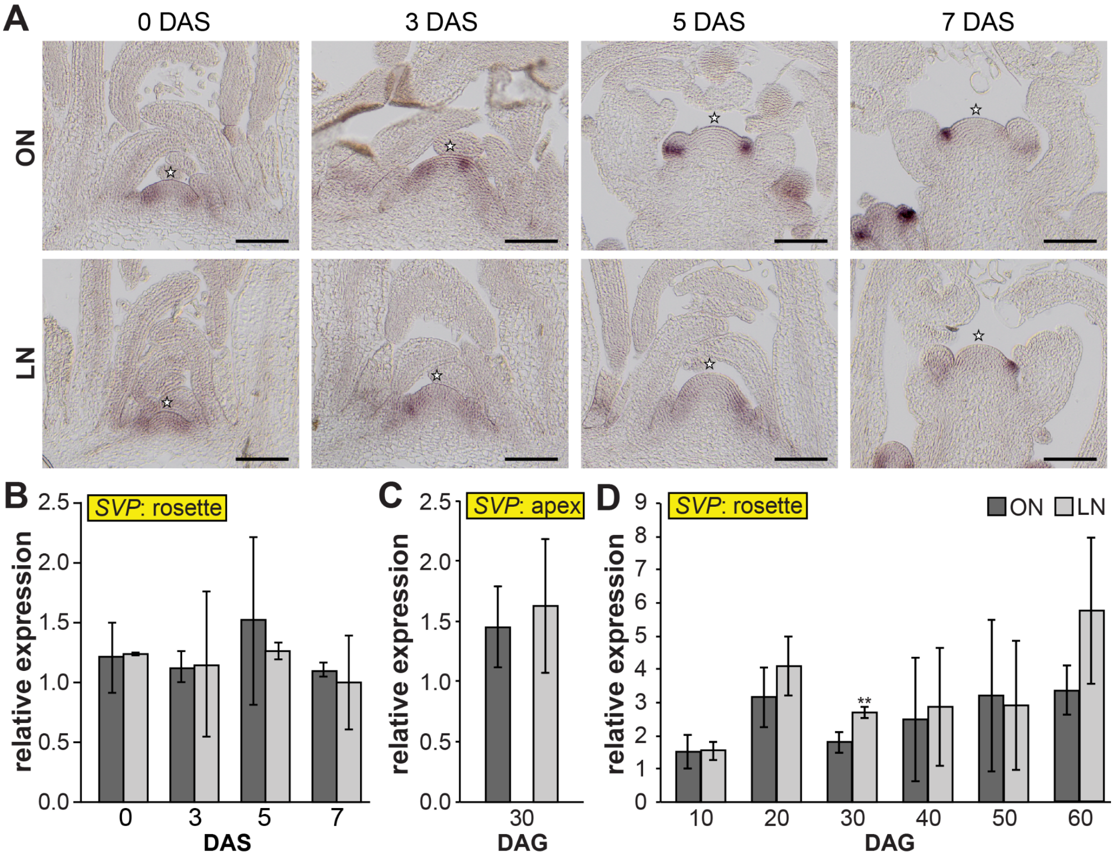


**Figure S5**: ***SHORT VEGETATIVE PHASE* in response to nitrogen limitation. (A)** RNA *in situ* hybridization using *SHORT VEGETATIVE PHASE* specific probe on longitudinal sections through apices of Col-0 plants grown in optimal nitrogen (ON) and limited-nitrogen (LN) soils under short days (8h light/ 16h darkness) for the first 30 days and then transferred to long days (16h light/ 8h darkness) to initiate the floral transition for 3, 5, and 7 days (SD-LD shift). **(B, C, D)** Expression measured by RT-qPCR in (B) apices of Col-0 plants grown in the SD-LD shift conditions, in (C) apices and (D) rosettes of plants continuously grown under short days. N = 3. Abbreviations: days after germination (DAG); days after shift (DAS). Data represents mean, error bars are standard deviations (s.d.), n=3, statistically significant difference between ON and LN (Student's *t*-test, $^{**}P<0.01$).

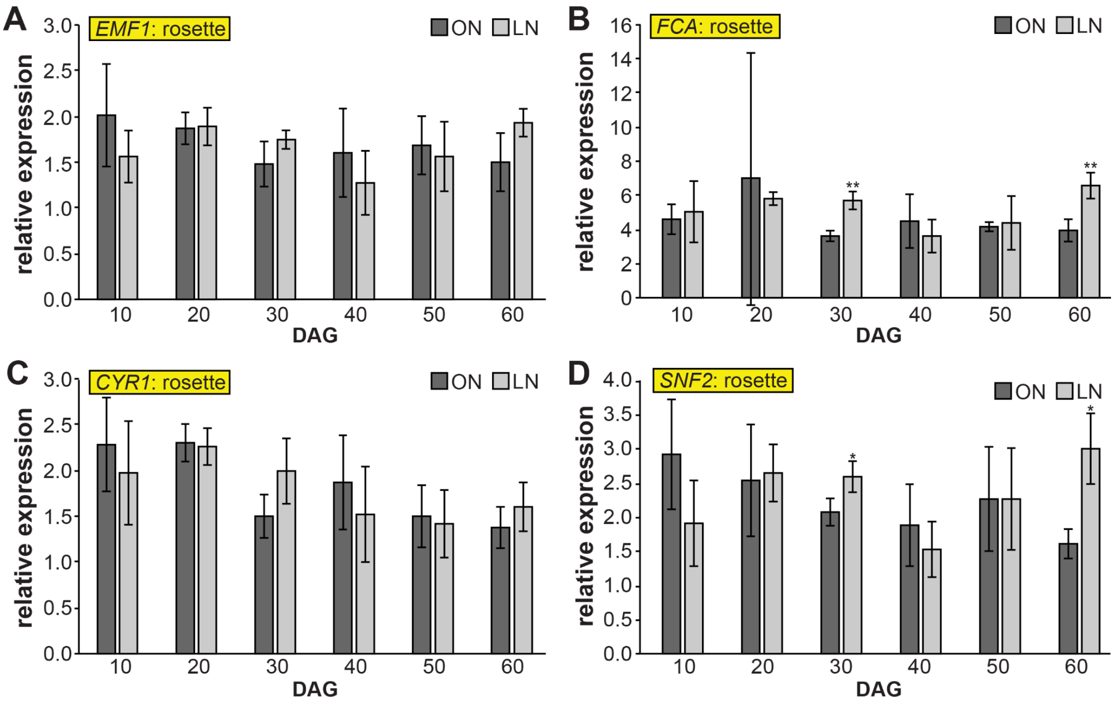


**Figure S6**: **Regulators upstream *FLOWERING LOCUS C* in response to N limitation.** (**A**, **B**, **C**, **D**) Expression of (A) EMF1, (B) FCA, (C) CRY1and (D) SNF2 measured by RT-qPCR in rosettes of Col-0 plants grown in optimal nitrogen (ON) and limited-nitrogen (LN) conditions under short days (16h light/ 8h dark). $n$ = 3. Data represents mean, error bars are standard deviations (s.d.), statistically significant difference between ON and LN (Student's *t*-test, *$P<0.05$, **$P<0.01$). Abbreviations: days after germination (DAG).

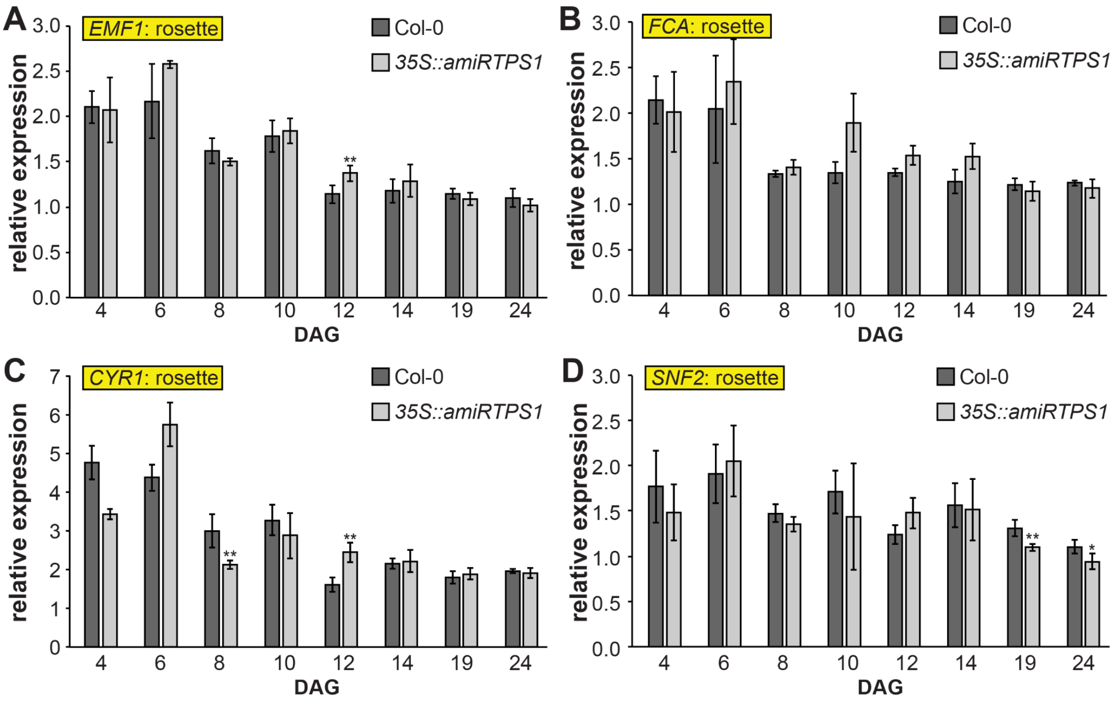


**Figure S7**: **Regulators upstream *FLOWERING LOCUS C* in *35S::amiRTPS1* plants.** (**A**, **B**, **C**, **D**) Expression of (A) EMF1, (B) FCA, (C) CRY1and (D) SNF2 measured by RT-qPCR in rosettes of Col-0 and *35S::amiRTPS1* plants grown under long days (16h light/ 8h darkness). *N* = 4. Data represents mean, error bars are standard deviations (s.d.), statistically significant difference compared to Col-0 wild-type (Student *t*-test, **P*<0.05). Abbreviations: days after germination (DAG).

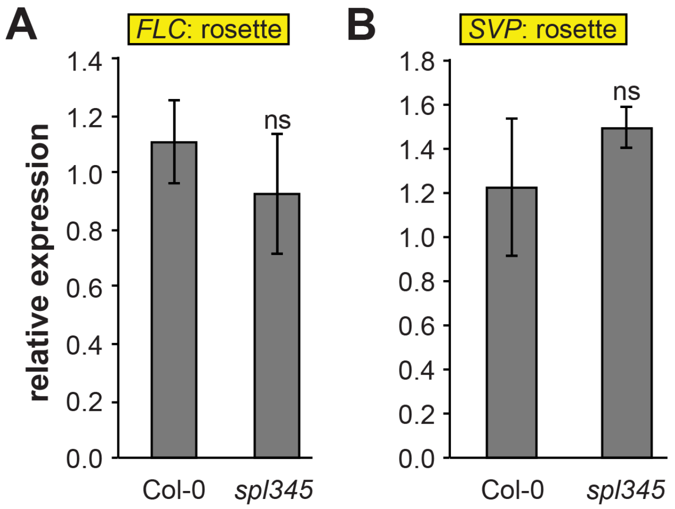


**Figure S8**: ***FLOWERING LOCUS C*** **and** ***SHORT VEGETATIVE PHASE*** **in** ***spl345*** **mutant plants.** (**A, B**) Expression of (A) *FLOWERING LOCUS C* and (B) *SHORT VEGETATIVE PHASE* measured by RT-qPCR at 20 days after germination in rosettes of wild-type Col-0 and *spl345* mutant plants plants grown under long days (16h light/ 8h darkness). *n* = 3. Data represents mean, error bars are standard deviations (s.d.). Statistical significance of the difference between genotypes was calculated by Student's *t*-test (ns – not significant).

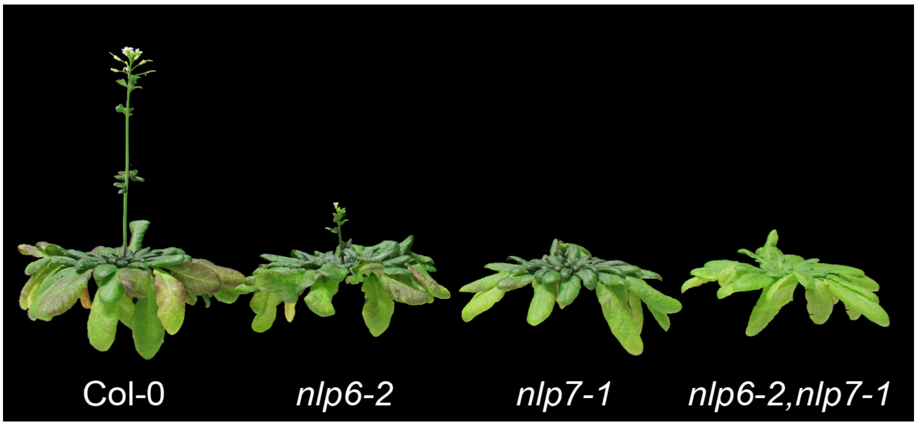


**Figure S9**: **Plant phenotype of *nlp6* and *nlp7* mutant plants.** Representative pictures of the plants analyzed in Figure 4 at 90 DAG.

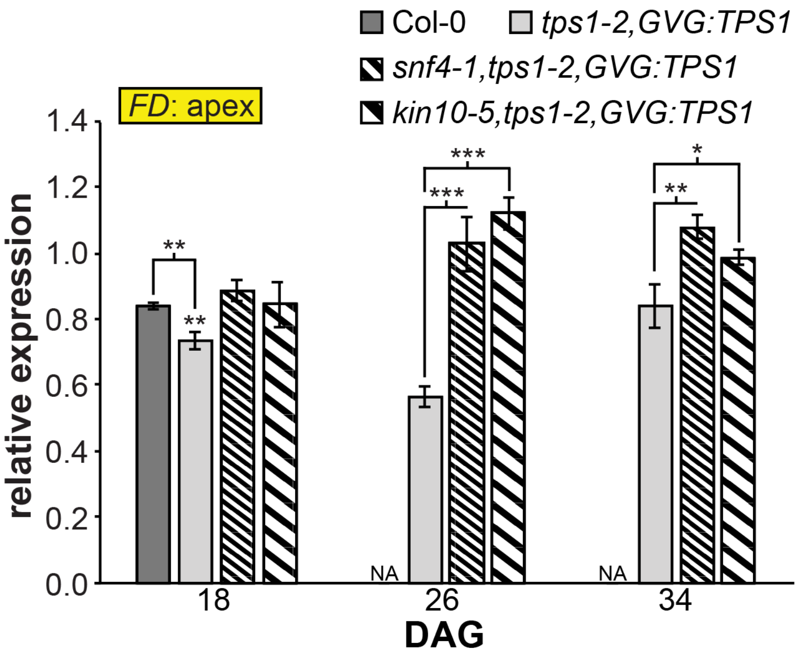


**Figure S10**: ***FLOWERING LOCUS D*** **in** ***snrk1,tps1-2,GVG::TPS1*** **mutants.** RNA-seq data from apices of Col-0, *tps1-2,GVG::TPS1, kin10-5,tps1-2,GVG::TPS1* and *snf4-1,tps1-2,GVG::TPS1* apices. n=3. Data represents mean of VST expression, error bars are standard deviations (s.d.), statistically significant difference compared to Col-0 wild-type (Student *t*-test, **P*<0.05, ***P*<0.01, ****P*<0.001). Data obtained from available online RNA-seq dataset (Zacharaki et al., 2022).

### Supplementary Tables

#### Table S1. Flowering time data of experiments described in this study.

Abbreviations: DTF, days to flowering (bolting time) in bold as referred to in the main text; TLN, total leaf number; n, number of individual plants; (+/-) standard deviation, presence or absence of significance based on Student's *t*-test calculated between plants grown on the ON and LN soils, respectively (+ : p-value < 0.05, - : p-value > 0.05).

| Plant lines/Experiment | ON | | | LN | | |
|---|---|---|---|---|---|---|
| | DTF | TLN | n | DTF | TLN | n |
| **Experiment 1 (long days) – Figure 1B** | | | | | | |
| Col-0 (wild type) | n.a. | **11.6 ± 0.9** | 20 | n.a. | n.a. | - |
| *flc-3* | n.a. | **9.5 ± 1.0** | 20 | n.a. | n.a. | - |
| *tps1-2, GVG::TPS1* | n.a. | **73.7 ± 4.6** | 20 | n.a. | n.a. | - |
| *flc-3, tps1-2, GVG::TPS1* | n.a. | **56.6 ± 5.1** | 20 | n.a. | n.a. | - |
| **Experiment 2 (short days/cold treated for 8 weeks) – Figure 3A** | | | | | | |
| Col-0 (wild type) | **89.3 ± 2.9** | 32.6 ± 2.3 | 17 | **87.9 ± 1.3**$^{(-)}$ | 30.5 ± 1.7 | 18 |
| *flc-3* | **91.2 ± 4.0** | 33.6 ± 4.3 | 18 | **92.4 ± 2.3**$^{(-)}$ | 35.0 ± 2.3 | 18 |
| **Experiment 3 (short days) – Figure 3B** | | | | | | |
| Col-0 (wild type) | **70.6 ± 4.5** | 68.3 ± 5.4 | 16 | **79.4 ± 2.4**$^{(+)}$ | 64.56 ± 2.0 | 16 |
| *flc-3* | **73.0 ± 5.1** | 65.3 ± 3.5 | 18 | **72.7 ± 2.9**$^{(-)}$ | 61.2 ± 3.2 | 18 |
| *svp-32* | **47.9 ± 3.1** | 33.9 ± 2.6 | 18 | **46.9 ± 2.5**$^{(-)}$ | 27.0 ± 2.2 | 18 |
| **Experiment 3b (short days) – repeat – Figure 3B** | | | | | | |
| Col-0 (wild type) | **66.2 ± 1.6** | 63.6 ± 2.3 | 16 | **81.1 ± 4.3**$^{(+)}$ | 62.1 ± 2.6 | 18 |
| *svp-32* | **44.1 ± 2.4** | 28.2 ± 2.1 | 19 | **46.3 ± 2.2**$^{(-)}$ | 28.1 ± 3.3 | 19 |
| **Experiment 3c (short days) – repeat – Figure 3B** | | | | | | |
| Col-0 (wild type) | **61.0 ± 2.4** | 61.4 ± 2.1 | 20 | **71.9 ± 2.1**$^{(+)}$ | 54.2 ± 2.0 | 17 |
| *flc-3* | **66.6 ± 3.0** | 59.5 ± 2.0 | 17 | **68.1 ± 2.2**$^{(-)}$ | 53.7 ± 2.3 | 18 |

### Table S2. Vegetative phase change data of experiment described in this study (Figure S2).

Abbreviations: JLN, juvenile leaf numbers in bold; n, number of individuals; +/- standard deviation, presence or absence of significance based on Student's *t*-test (+ : p-value < 0.05).

| Plant line | JLN | n |
|---|---|---|
| Col-0 (wild type) | **4.7 ± 0.5** | **20** |
| *tps1-2,GVG::TPS1* | **9.8 ± 0.9$^{(+)}$** | **20** |
| *flc-3* | **5.0 ± 0.7$^{(+)}$** | **20** |
| *flc-3,tps1-2,GVG::TPS1* | **8.2 ± 1.0$^{(+)}$** | **20** |

**Table S3. Putative nitrate responsive *cis*-elements (NREs) in regulators upstream *FLOWERING LOCUS C*.**

| Gene | Locus ID | NRE |
|---|---|---|
| NIR1 (consensus NRE) | AT2G15620 | tGaCCctT---(n)---AAGaG |
| FLC | At5g10140 | - |
| FRI | At4g00650 | TGACCgaTcatAAGAGAAGAG<br>TGACCaTTgatatatttatttctcaacagaaagAAGcG<br>aGACCacTaataagataccAAGtG<br>aGcCCaTTaccAAGAG |
| SUF4 | At1g30970 | - |
| ELF7 | At1g79730 | - |
| PIE1/SNF2 | At3g12810 | cGaCCtTTccttctacggcgcctaaaAAGgG |
| SEF | At5g37055 | - |
| VIN3 | At5g57380 | - |
| VRN1 | At3g18990 | - |
| VRN2 | At4g16845 | - |
| EMF1 | At5g11530 | TGaCCgTTttcagAAGAG |
| EMF2 | At5g51230 | TGtCCaTTgctgcagctaaagtccatgagtgaggaaAAGt<br>G |
| TFL2 | At5g17690 | - |
| FCA | At4g16280 | cGtCCaaTgggtcctaacggtggtgtgggaggagAAGgG |
| FVE | AT2G19520 | - |
| HUA2 | AT2G19520 | - |

### Table S4. Oligonucleotides used in this study.

| Gene (AGI) | Oligo | Sequence (5'>3') | Product lengths (bp) |
|---|---|---|---|
| **Oligonucleotides used for qRT-PCR** | | | |
| ***TUB2*** | P-344 | GAGCCTTACAACGCTACTCTGTCTGTC | 167 |
| At5g62690 | P-345 | ACACCAGACATAGTAGCAGAAATCAAG | |
| ***SAND*** | P-346 | AACTCTATGCAGCATTTGATCCACT | 61 |
| At2g28390 | P-347 | TGATTGCATATCTTTATCGCCATC | |
| ***UBI10*** | P-348 | CACACTCCACTTGGTCTTGCGT | 71 |
| At4g05320 | P-349 | TGGTCTTTCCGGTGAGAGTCTTCA | |
| ***PDF2*** | P-350 | TAACGTGGCCAAAATGATG | 61 |
| At1g13320 | P-351 | GTTCTCCACAACCGCTTGGT | |
| ***FLC*** | P-402 | GAAGACCGAACTCATGTTGAAGCT | 114 |
| At5g10140 | P-403 | GCTCCCACATGATGATTATTCTCC | |
| ***SOC1*** | P-532 | TTGAGCAGCTCAAGCAAAAGGA | 68 |
| At2g45660 | P-533 | TCCCCACTTTTCAGAGAGCTTCTC | |
| ***SPL3*** | P-544 | GAGTTTGTCAGGTCGAGAGTTGTACC | 74 |
| At2g33810 | P-545 | GCAGACTTTGTGTCGTTTGTGGT | |
| ***SPL4*** | P-546 | AATGGTCAGGTGGTGATGCAG | 61 |
| At1g53160 | P-547 | GCATAGGAAGTGTCATCTCTACCCTT | |
| ***SPL5*** | P-548 | CAGCAGGTTTCATGAGCTACCAG | 107 |
| At3g15270 | P-549 | CAAAACTGTCACCAGAGATCTTCCTC | |
| ***SVP*** | P-556 | CGGAGTCTATTACTAACGCCGGA | 89 |
| At2g22540 | P-557 | ATACGGTAAGCCGAGCCTAAGG | |
| **Oligonucleotides used for genotyping** | | | |
| ***FLC*** | P-0700 | ATGGGAAGAAAAAAACTAGAAATC | |
| At5g10140 | P-0701 | CTAATTAAGTAGTGGGAGAGTCAC | |
